# Dia1 Coordinates Differentiation and Cell Sorting in a Stratified Epithelium

**DOI:** 10.1101/2021.01.04.425231

**Authors:** Robert M. Harmon, John Devany, Margaret L. Gardel

## Abstract

Although implicated in adhesion, few studies address how actin assembly factors guide cell positioning in multicellular tissue. The formin, Dia1, localizes to the proliferative basal layer of epidermis. In organotypic cultures, Dia1 depletion reduced basal cell density and resulted in stratified tissue with disorganized differentiation and proliferative markers. Since crowding induces differentiation in epidermal tissue, we hypothesized that Dia1 allows cells to reach densities amenable to differentiation prior to stratification. Consistent with this hypothesis, forced crowding of Dia1-deficient cells rescued transcriptional abnormalities. Dia1 promotes rapid growth of lateral adhesions, a behavior consistent with the ability of cells to remain monolayered when crowded. In aggregation assays, cells sorted into distinct layers based on Dia1 expression status. These results suggested that as basal cells proliferate, reintegration and packing of Dia1-positive daughter cells is favored while Dia1-negative cells tend to delaminate to a suprabasal compartment. These data demonstrate how formin expression patterns play a crucial role in constructing distinct domains within stratified epithelia.

**Summary:** Harmon et al demonstrate that differential expression of an actin nucleator, the formin, Dia1, drives cell sorting and maintains distinct morphological domains within an epithelial tissue. This illuminates the possible utility of evolving a large formin family in orchestrating the compartmentalization and differentiation of complex tissues.

## Introduction

Throughout adulthood, epithelia turn over yet retain distinctive architectures. Such tissues often encompass subdomains distinguished by unique cellular morphology and behavior. The mechanisms, however, by which these domains arise and persist remain a puzzle. Exemplifying the problem, keratinocytes establish stratified structures at the skin surface characterized by layers with distinct morphologies (Luxenburg and Zaidel-Bar, 2019; Simpson et al., 2011). A basal layer of proliferative, columnar cells pack atop a collagenous dermis and give rise to overlying layers with flattened, squamous morphologies. As cells stratify and change shape, a differentiation program ensures that suprabasal cells repress proliferative behavior and express factors required for producing an effective barrier to water loss, pathogens and mechanical insult (Barrandon and Green, 1985).

Of note, basal cell shape affects the capacity to coordinate stratification with differentiation (Box et al., 2019; Miroshnikova et al., 2017). Reduced cell spread area, concomitant with basal layer packing, also triggers serum-response factor (SRF), YAP and NOTCH responses which drive differentiation (Connelly et al., 2010; Totaro et al., 2017). Coordination of cell movement with differentiation in these models, however, is contingent upon establishing a densely packed basal layer. The question arises as to how the skin and other stratified epithelia shunt proliferation into packing the basal layer rather than distributing cells homogenously throughout the tissue thickness. To compartmentalize tissues, cells sort out based on differential intercellular adhesion and actomyosin contractility (Fagotto, 2014; Maitre et al., 2012; Manning et al., 2010). The segregation of like cells based on these properties guides prominent developmental processes like gastrulation but is also thought to aid in maintaining compartments at homeostasis (Krens and Heisenberg, 2011). In epidermal tissue, genetic studies indicate that cadherin and integrin-based adhesion ensure proper basal layer formation (Brakebusch et al., 2000; Raghavan et al., 2000; Tinkle et al., 2008; Vasioukhin et al., 2001). Likewise, classic *in vitro* studies indicate that when mixed in a cell aggregate, undifferentiated keratinocytes separate from differentiated neighbors to form compartments (Watt, 1984). Recent studies highlight differences in basal and suprabasal layer stiffness (Fiore et al., 2020), pointing to mechanics as a likely determinant of this separation.

How actin regulatory proteins, through their effects on cell-cell adhesion and cortical mechanics, enforce compartmentalization of the basal layer is not well understood. Whereas prior studies implicate depolymerizing agents like cofilin in epidermal function (Luxenburg et al., 2015), the role of actin nucleators requires further attention. Disruption of the branched actin nucleator, Arp2/3, or its activator, the WAVE complex, perturbs epidermal function with varying effects on structure (Cohen et al., 2019; van der Kammen et al., 2017; Zhou et al., 2013). Unbranched actin filaments produced by the formin family anchor adhesions and participate in physiological adaptations to mechanical stretch within the epidermis (Aragona et al., 2020; Kobielak et al., 2004). Endogenous miRNA restricts expression of the canonical formin, Dia1, to the basal layer of human epidermis and tongue (Sundaram et al., 2013; Sundaram et al., 2017). Though diabetic ulceration and certain carcinomas yield aberrant expression patterns (Chakraborty et al., 2010; Sundaram et al., 2013; Sundaram et al., 2017; Whitson et al., 2018; Xing and Liu, 2017; Yang et al., 2019), the normal physiological function of Dia1 in the basal layer remains unclear. One hypothesis is that differential Dia1 expression translates into differential adhesion effects amenable to packing and producing a properly shaped basal compartment.

In addressing this hypothesis, we found that Dia1 depletion in a model epidermis disrupted basal layer morphology and led to differentiation defects. Effects on morphology originated from the capacity of Dia1 to accelerate lateral adhesion growth. In monolayers, this feature allowed cells to accommodate crowding by enabling redistribution of the cortex towards building taller lateral adhesions and smaller basal footprints. Dia1 depletion, in contrast, forced cells to evade crowding via delamination. Thus, as proliferation added cells to the Dia1-positive basal layer, the ensuing rearrangement of junctions favored packing the basal layer over movement into the suprabasal compartment. This effect ensured a density amenable to triggering differentiation. *In* vitro assays also supported the hypothesis that Dia1 expression discrepancies can guide cell compartmentalization. In mature stratified epithelia, we posit that the basally skewed expression pattern of Dia1 establishes a differential adhesion effect which guides redistribution of adhesions during division. Rapid expansion of lateral junctions between Dia1-positive neighbors ensures reintegration of basal daughter cells during division. In contrast, crowding of a Dia1-negative cell by a neighboring division drives delamination and stratification. Our results provide the first indication that formin expression gradients, through effects on cell sorting, enforce the maintenance of distinct morphological zones within complex tissues.

## Results

### Dia1 controls basal layer morphology and packing

Involvement of Dia1 in immune function and the sensitivity of skin to immunological influence steered us toward the *in vitro* approach of a well characterized, organotypic culture model (Albanesi et al., 2018; Lin et al., 2011; Peng et al., 2007). To mimic the *in vivo* juxtaposition of epidermal epithelia to its supporting connective tissue, three-dimensional cultures are produced by plating keratinocytes, like the HaCaT line, upon fibroblast-doped collagen (Schoop et al., 1999). Importantly, cultures developed at an air-liquid interface construct a stratified tissue with architecture and expression profiles mimicking those of normal tissue (Koehler et al., 2011; Sundaram et al., 2017). Immunostaining of tissue sections produced from 10 day old, paraffin embedded control (CTL) cultures indicated that Dia1 expression, as well as the shape and orientation of nuclei, changed with respect to position in the stratified tissue. Dia1 concentrated in the basal layer, approximated as keratinocytes lying within 10 μm of the underlying collagen (dashed line, Fig. 1A+B). Basal layer keratinocytes produced a mean linear density of 12 cells per 100 μm (Fig. 1C). Within the basal layer, nuclei are elongated, with an average aspect ratio of 2.5 (Fig. 1E), and are oriented such that the long axis is nearly perpendicular to the basal surface, with the average angle between these axes being 20 degrees (Fig. 1B+D). Both the nuclear shape and orientation change in a stereotypical fashion towards the apical surface. As their relative position increases, the nuclei are less elongated but maintain their orientation (Fig. 1A-E). In the suprabasal layers, the aspect ratio sharply increases and orientation increases such that the long axes are parallel to the basal interface (Fig. 1A-E).

**Figure 1.**
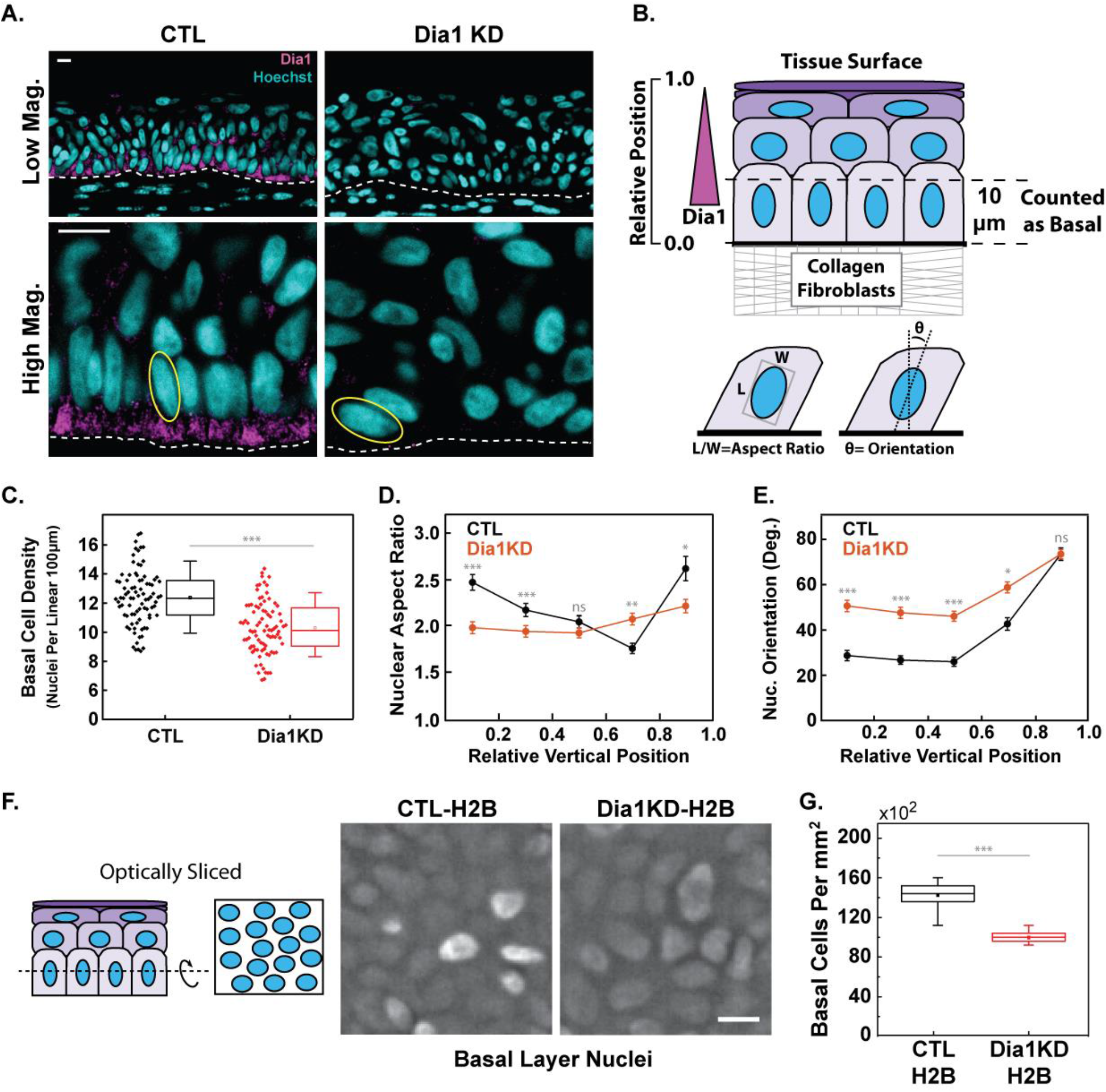
Dia1 Supports Basal Layer Packing In Stratified Tissue. A) Immunohistochemical staining of Dia1 in cross-sections obtained from paraffin embedded cultures, fixed 10 days after lifting to an air-liquid interface. Scale bars=10μm. Example of fit ellipses used for subsequent measurements indicated in yellow. Dotted line separates keratinocytes from the underlying collagen and fibroblasts. B) Schematic illustrating measurements taken from tissue cross-sections and presented in panels C-E. C) Linear density of basal nuclei, 85-87 images pooled from two expts. D) Aspect ratio of nuclei as a proxy for cell shape throughout tissue thickness. E) Orientation of nuclei throughout the tissue thickness with respect to the keratinocyte-collagen border. For panels D and E, plotted points represent the mean +/− se of data lying within bins bounded by x-axis ticks. F) Optical sectioning of basal nuclei in live 3D cultures carrying GFP-labeled H2B, 6 days after lifting to an air-liquid interface. Images are average projections of confocal slices extending 6μm above the most basal, in-focus plane of H2B signal and filtered to remove background. Scale bar= 10μm. G) Density of basal nuclei obtained from optical sectioning. Data represent 11 fields pooled from two independent experiments. For panels C and G, Boxes=25-75th percentile, whiskers= 10-90th percentile. Student’s t-test results are indicated as: p<0.05 (*), p<0.01(**), p<0.0001(***), p>0.05 was considered not significant (ns).

To explore the role of Dia1 on the architecture of stratified tissue, we used CRISPR and shRNA to reduce Dia1 expression (Fig. S1). Both techniques diminished Dia1 expression, with shRNA yielding knockdown efficiencies of 65-80% and CRISPR-edits causing near total abrogation of Dia1 protein levels. Next generation sequencing identified two frameshift variants with premature stop codons, accounting for ~85% of the genomic sequencing signal (Fig. S1). Tissue depleted of Dia1 by CRISPR (Dia1KD) stratified but produced a basal layer that was 20% less dense than CTL cells and lacked the stereotypical, columnar morphology (Fig. 1A, C). Within the basal layer, the nuclei of Dia1KD cells were less elongated and randomly oriented, at times lying parallel to the underlying collagen (Fig. 1A, D, E). The nuclear shape remained constant in the suprabasal layers and showed a similar trend towards becoming parallel to the basal interface at high relative positions (Fig. 1E). Multicellular patches of Dia1KD cells were also observed protruding into the underlying collagen (Fig. S1).

To overcome the limitations of measuring 2D density in cross-section, we conducted confocal imaging of flat mounted, H2B-labeled cells (CTL-H2B and Dia1KD-H2B) cultured for 6 days (Fig. 1F). These data revealed similar basal layer density deficits in Dia1KD cell lines (Fig. 1G). Dia1-depleted tissue successfully stratified but did so without producing a properly packed basal layer, leading us to posit that cells were escaping the basal layer before reaching an appropriate density.

### Dia1 Expression Supports Epidermal Differentiation

Since morphology dictates the success with which basal cells differentiate (Connelly et al., 2010; Totaro et al., 2017), we next tested whether the aberrant morphology caused by Dia1 depletion impacted epidermal differentiation. To address this possibility, sections of organotypic cultures were immunohistochemically stained for the differentiation marker, keratin-10 (KRT10). As in normal tissue, CTL cells produced a three-dimensional culture in which KRT10 localized specifically to suprabasal layers, comprising 60% of the cross-sectioned, epidermal area (Fig. 2A+B). Dia1KD cells, conversely, produced tissue that stratified but contained lesions of KRT10-negative cells (Fig. 2A+B). Organotypic cultures derived from shRNA-treated cells (shCTL and shDia1) corroborated these results (Fig. S2).

**Figure 2.**
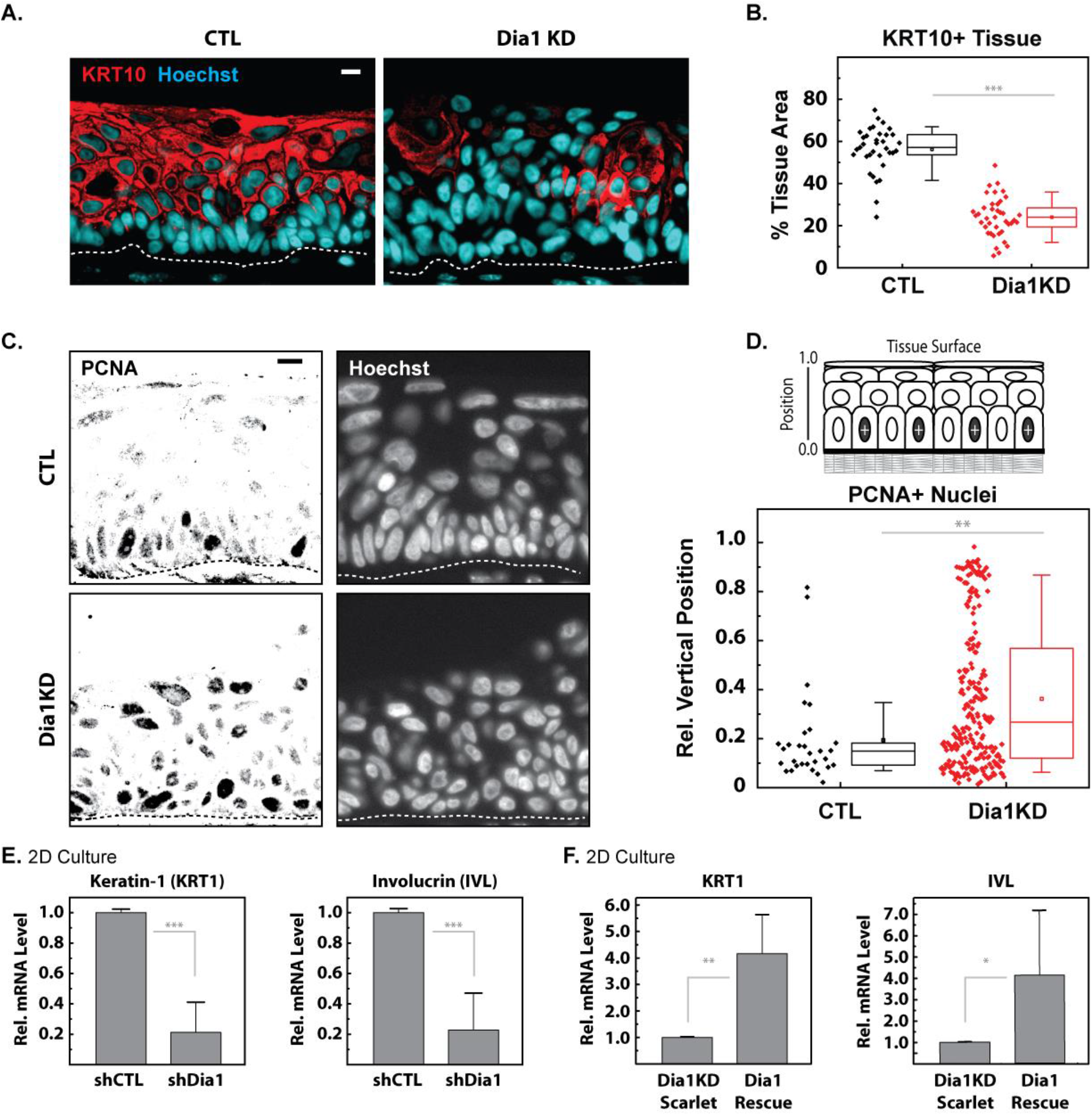
Dia1 supports epidermal differentiation. (A) Keratin-10 (KRT10) staining of cross-sectioned, paraffin embedded 3D cultures. B) Percent cross-sectional area expressing KRT10 (3-fold greater signal than mean acquired from control basal layers). Represents 35-40 images per condition pooled from two experiments. C) PCNA staining of paraffin sections. D) Distribution of PCNA+ cells with respect to vertical position in the epidermis. PCNA+ = 1.5 fold greater signal than control, suprabasal nuclei. Respectively, 1036 and 1324 nuclei were analyzed from control and Dia1-deficient tissue. Data was pooled from two experiments, representing 40 images per condition. E) Effects of shRNA-mediated Dia1 depletion on differentiation marker expression 48hrs post calcium addition in 2D culture. Results represent 6-9 technical qPCR replicates pooled from two (KRT1) to three (IVL) experiments. +/− SD. F) Differentiation induction at 24hrs by ectopic mDia1 introduced to Dia1KD cells. Results pooled from 6 technical replicates obtained from two experiments. +/−SD. For A) and B), scale bars=10μm; dashed line = approximate collagen boundary. For panels C) and D), boxes=25-75th percentile, whiskers= 10-90th percentile, square= mean, line = median. Student’s t-test results are indicated as: p<0.05 (*), p<0.01(**), p<0.0001(***).

Spatially-segregated proliferation is an important attribute of differentiation in stratified tissue, its loss linked to pathologies ranging from barrier disruption to viral oncogenesis (Moody and Laimins, 2010; Rorke et al., 2015). Staining for the proliferation marker, PCNA, allowed for visualization of where proliferative cells reside in the stratified layers. Control tissue demonstrated proper restriction of proliferative cells to the basal layer (Fig. 2C+D). Dia1-depletion led to the extension of PCNA positive cells throughout the tissue thickness (Fig. 2C+D). Further, the percentage of PCNA positive nuclei was 18% in the Dia1KD cells, reflecting a six-fold overall increase from that of control tissue (2.9%). This is reminiscent of hyperproliferative phenotypes associated with stratified tissue disorders like psoriasis and esophageal disease (Hugh and Weinberg, 2018; Wang et al., 2005).

Standard two-dimensional culture methods of assessing differentiation corroborated the organotypic model results. A well-known technique for inducing differentiation *in vitro* entails culturing cells in low calcium (0.03-0.07mM CaCl_2_), then, switching the cells to a high calcium (1.2-2.8mM) media (Deyrieux and Wilson, 2007). Compared to shCTL-treated cells, the mRNA level of differentiation markers involucrin (IVL) and keratin-1 (KRT1) were diminished 5-fold in shDia1 cells (Fig. 2E). To validate the specificity of this result, we transduced Dia1KD cells with a Scarlet-tagged, full length mouse Dia1, referred to hereafter as Dia1-Rescue cells (Fig. S2). Compared to Scarlet-only controls (Dia1KD-Scarlet), reintroduction of Dia1 facilitated induction of IVL and KRT1 expression in this simplified assay (Fig. 2F). These results confirmed the specificity of differentiation abnormalities stemming from Dia1 depletion.

### Forced Crowding Rescues Transcriptional Abnormalities in Dia1-deficient Cells

To assess the impact on transcriptional changes expected during stratification, we assessed differential gene expression in Dia1KD-Scarlet cells compared to the Dia1-Rescue line via RNAseq. Of the 15,150 genes detected, approximately 275 of these were upregulated at least 2-fold in the rescue line (Fig. 3A). Strikingly, the top five gene sets enriched by Dia1 rescue, as detected by automated gene ontology, were those of cornification, keratinization, keratinocyte differentiation and epidermal differentiation/development. Although highly overlapping sets, these strongly implicated Dia1 in epidermal development.

**Figure 3.**
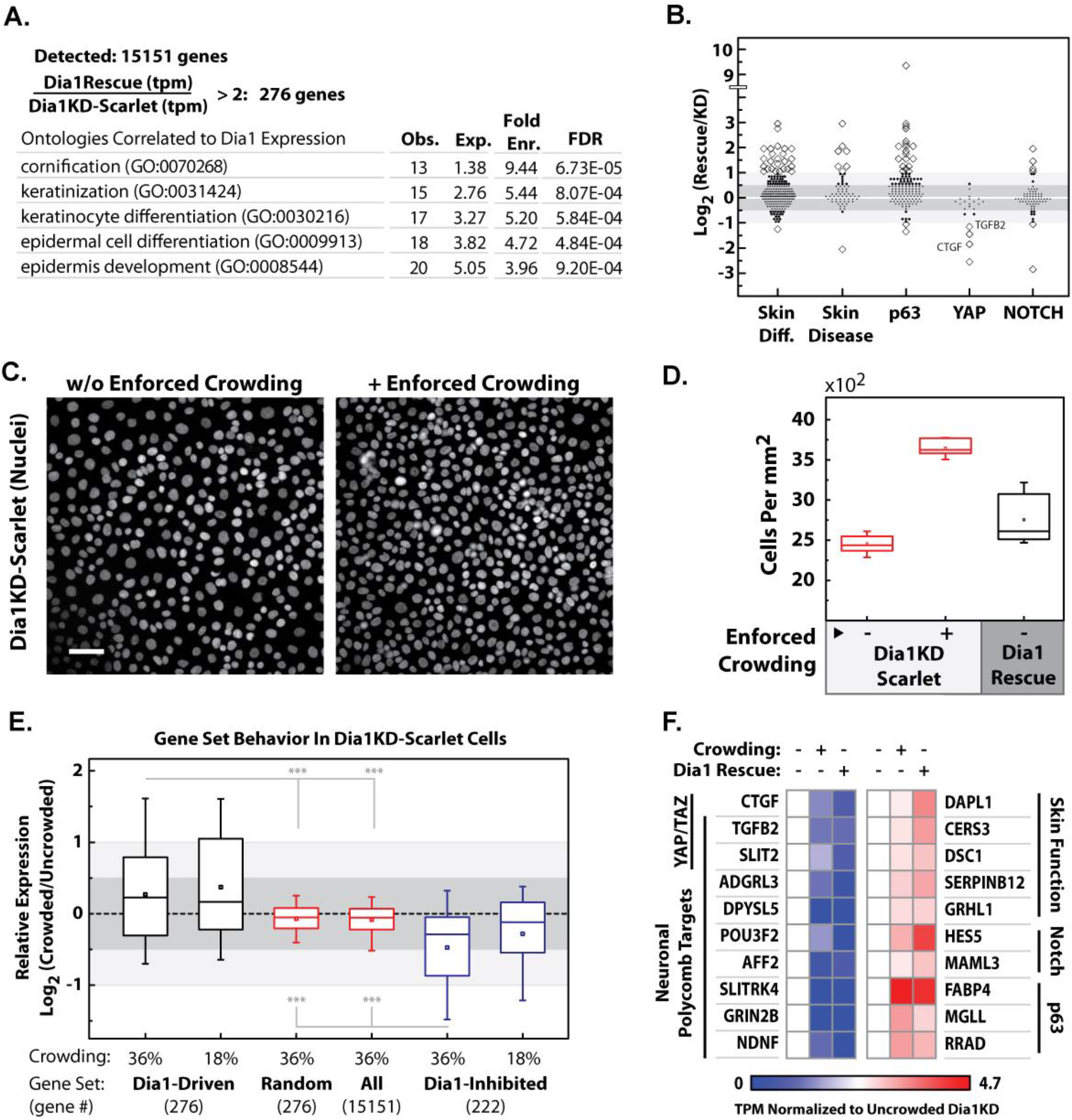
Crowding rectifies transcriptional defects associated with Dia1-deficiency. A) Gene ontology analysis of genes identified by RNAseq as being upregulated >2-fold in response to ectopically expressed mDia1 in Dia1KD cells. B) Log2 fold change analysis of Dia1KD-Scarlet cultures compared to Dia1-Rescue cells with respect to published gene sets tracking differentiation, monogenic skin disorders, genes induced by p63, YAP/TAZ and NOTCH signaling. Prominent YAP targets are noted. C) Cells plated on biaxially stretched PDMS (36% by area) were released from stretch to force cell crowding. Images depict nuclear staining of uncompressed and compressed cells immediately after releasing stretch. Parallel cultures were incubated a further 5hrs and processed for RNAseq. D) Crowding quantified in terms of cell density with and without forced crowding.30 fields per condition. E) Behavior of genes identified as Dia1-responsive (induced or inhibited >2-fold by Dia1 rescue) within Dia1KD-Scarlet cells subjected to crowding. F) Effect of crowding versus Dia1 rescue on genes belonging to salient signaling groups within Dia1KD-Scarlet cells. Expression analyses are based on TPM (transcripts per million) normalizations. Scale bars= 50μm. Box plots=25-75 percentile, whiskers=10-90th percentile, line= median, square= mean. T-test results of p<0.0001(***) are indicated.

We then performed a manual query of this data with published gene sets for skin differentiation (Joost et al., 2016), skin disease (Lemke et al., 2014) and genes driven by salient pathways like p63 (Truong et al., 2006), YAP (Wang et al., 2018) or NOTCH signaling (Wang et al., 2014). Genes linked to skin disease, differentiation, p63, or NOTCH signaling increased in response to Dia1 rescue (Fig. 3B). In particular, we observed increased expression of the small proline-rich (SPRR) proteins (Fig. S3), essential for cornification (Carregaro et al., 2013). In contrast, several notable YAP target genes decreased in response to rescued Dia1 expression. Targets of other signaling programs salient to differentiation like SRF (Olson and Nordheim, 2010), SHH (Lewandowski et al., 2015) and Wnt (Watanabe et al., 2014) did not demonstrably trend towards up- or down-regulation, but contained individual transcripts displaying sensitivity to Dia1 expression (Fig. 3B, Fig. S3). Altogether, this data underscored that Dia1 expression, even in *in vitro* culture models, could drive gene expression signatures pertinent to *in vivo* epidermal differentiation.

To determine whether aberrant cell area and/or density contributed to the differentiation phenotype of Dia1-deficient cells, we explored whether enforced crowding could rescue transcriptional abnormalities. Inspired by a previous study (Miroshnikova et al., 2017), keratinocytes were seeded on biaxially pre-stretched (36% by area) PDMS sheets. After incubating cells for 20hrs, stretch was released, in effect, increasing cell density. Sampled at the center of the culture area, release from stretch increased Dia1KD-Scarlet density by ~50% from 2455 to 3645 cells per mm^2^ (Fig. 3D). By comparison, a parallel Dia1-Rescue culture produced a mean cell density of 2755 cells per mm^2^ (Fig. 3D). Thus, relaxing pre-stretch provided a means to enforce crowding of Dia1-deficient cells into densities beyond those achieved by unmanipulated Dia1-Rescue cells. Mitigating the pre-stretch to 18% and, in turn, dampening the extent of cell density increase, enabled achievement of densities more closely mimicking those of Dia1-Rescue cultures. Following 5hrs of incubation post-release, samples were collected for RNA sequencing (Fig. S2). Dia1KD-Scarlet cells with and without enforced crowding were compared to identify differentially expressed genes. Interestingly, we saw enhanced expression of the 276 Dia1-driven genes identified in Fig. 3A for both levels of enforced crowding. Similarly, we saw reduced expression of Dia1-Inhibited genes. This appeared specific to Dia1-responsive genes, as randomly chosen gene sets demonstrated little response to crowding (Fig. 3E).

Manual inspection of transcripts from Fig. 3E that were at least 1.6x enriched or depleted identified a number of differentiation markers as well as components of the pro-differentiation p63 and Notch pathways (Fig. 3F). Strikingly, GSEA database analysis revealed that both Dia1 expression and enforced crowding of Dia1KD-Scarlet cells suppressed non-epidermal (neural; GO:0007417, GO:0022008) gene expression, specifically amongst targets of Polycomb Repressive Complex 2 (PRC2)-mediated silencing (Ben-Porath et al., 2008). This set included two YAP-activated genes, TGFB2 and SLIT2, which tracked with CTGF in responding negatively to crowding (Cordenonsi et al., 2011; Wang et al., 2018). Consistent with reports of YAP antagonizing differentiation (Totaro et al., 2017; Zhou et al., 2013), both crowding and Dia1 rescue upregulated expression of differentiation markers including DAPL1, DSC1 and CERS3. Since mechanical stimulation elicits a PRC2 response in keratinocytes (Le et al., 2016) and abrogated epidermal identity defaults to a neural signature in p63-deficient cells (De Rosa et al., 2009; Mardaryev et al., 2015), we posited that Dia1-dependent crowding supported differentiation, in part, by protecting the more basic attribute of epidermal identity. Together, the YAP and PRC signatures suggested that Dia1-dependent effects on basal layer crowding contributed to keratinocyte differentiation.

### Dia1 Expression Controls the Density At Which Cells Stratify

To explore the formation of stratified tissue, we used live cell imaging to monitor the onset of suprabasal layer production in cultures of GFP-H2B marked cells plated atop fluorescently labeled collagen (Fig. 4A). CTL-H2B and Dia1KD-H2B cells initially formed monolayers (Fig. 4A). At 24 hrs, basal layer density in CTL-H2B cultures increased by 2-fold, accompanied by the onset of a suprabasal layer (Fig. 4A+B). Over the next 75 hrs, the basal layer density remained constant while the number of suprabasal cells continued to increase (Fig. 4A+B). In comparison, Dia1KD-H2B cells populated the culture at a reduced rate, possibly due to slower rates of proliferation (Fig. 4B; Fig. S4). The basal layer, nonetheless, did achieve a steady state density around 24 hrs post plating, albeit 20% lower than that of CTL-H2B cells (Fig. 4B). Though cells were observed in the subrabasal layer, the basal layer density was significantly reduced at all times. To measure the basal layer density corresponding to the onset of suprabasal layer formation, we measured the linear cell density in regions proximal to instances with 1-3 suprabasal cells over a 160 μm slice. Consistent with our earlier results, the linear density of the basal cell layer was 20% reduced in Dia1KD cells compared to the control cells. Thus, Dia1-deficient cultures successfully stratified but did so later and with lower basal cell densities (Fig. 4C).

**Figure 4.**
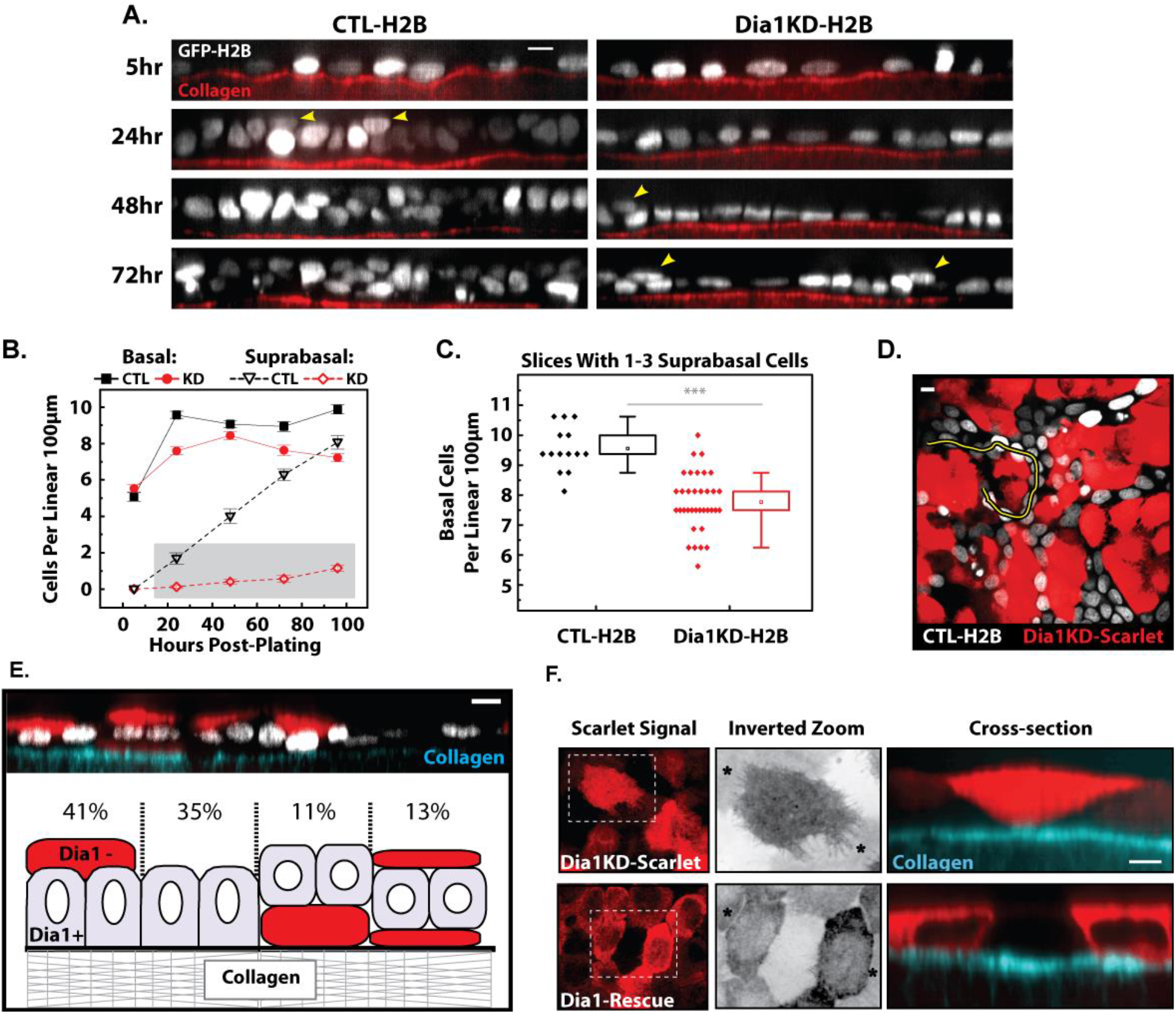
Dia1 expression controls the density at which cells protrude and migrate atop neighbors. A) Orthogonal slices, averaged across 5μm thicknesses, of live, GFP-H2B marked cells, plated upon labeled collagen gels for the indicated times. Yellow arrowheads = suprabasal cells. B) Counting of basal and suprabasal cells per orthogonal slice with respect to time since plating. 20 orthogonal slices per condition per timepoint were analyzed. H2B signals occluding >50% or an underlying H2B signal or overlying multiple H2B-labeled nuclei were considered suprabasal. Plotted values represent the mean +/− se per slice. Gray region: data pertaining to panel C. C) Linear density of basal cells in z-slices, limited to slices containing only 1-3 suprabasal cells, regardless of timepoint. Diamonds= individual slices. Boxes= 25-75th percentile, whiskers = 10-90th percentile, square = mean. D) Mixed cultures containing CTL-H2B cells (white) and Dia1KD-Scarlet cells (red) at 96hrs post plating. Image of basal layer cells. Yellow line pertains to panel E. E) Orthogonal slice along CTL-H2B nuclei bordering a Dia1KD-Scarlet region marked by yellow in panel D. Frequencies of illustrated spatial relationships between CTL-H2B and Dia1KD-Scarlet cells were determined for 20 boundary regions encompassing 180 control nuclei. F) Maximum projections of the top 2μm of confocal slices capturing Scarlet and Scarlet-tagged mDia1 in Dia1KD-Scarlet and Dia1-Rescue cells, respectively. Inverted zoom depicts boxed regions in left panels, cross-sections were taken between the asterix marks in the inverted panels. Scalebars= 10μm for panels A, D and E; 5μm for F. Student’s t-test results are indicated where p<0.0001(***).

Dia1 expression is sharply attenuated in the suprabasal cells of control tissue. We wished to explore whether this segregation would spontaneously occur in mixed cultures of Dia1KD and CTL cells. We produced mixed cultures containing even portions of Dia1KD cells expressing the fluorescent protein Scarlet, marking the cytoplasm, with CTL-H2B cells. After 4 days of growth on collagen gels, confocal imaging was used to assess the spatial relationship between Dia1KD-Scarlet and CTL-H2B cells (Fig. 4D). The two cell types did not readily mix; CTL-H2B cells formed strands of connected islands weaving in between Dia1-negative regions (Fig. 4D). By assessing orthogonal sections taken from control cells bordering Dia1-negative domains, we noted that 35% of control cells lacked suprabasal partners (Fig. 4E). This behavior was distinct from homogenous control cultures within which all basal cells had suprabasal neighbors at this timepoint (Fig. 4A+B), indicating that the presence of Dia1KD-Scarlet cells hampered CTL-H2B stratification. Nonetheless, we found that 41% of CTL-H2B cells had acquired Dia1KD-Scarlet cells as suprabasal neighbors whereas only 11% of CTL-H2B cells had assumed a suprabasal position overlying a Dia1KD-Scarlet cell (Fig. 4E). The remaining 13% contained CTL-H2B cells wedged between two layers of Dia1KD-Scarlet cells (Fig. 4E).

To understand how this sorting occurred, we then considered how Dia1 expression impacted the cross-sectional shape of cells, as a means of inferring the likelihood of apical protrusion. A previous study noted that Diaphanous depletion in Drosophila led to enhanced apical cross-sections, consistent with increased apical protrusion (Homem and Peifer, 2008). To facilitate this measurement, we formed monolayers and exploited the cytoplasm outlines produced by Scarlet and Scarlet-mDia1 signals in Dia1KD-Scarlet and Dia1-Rescue cells, respectively. Immunofluorescence of fixed cells revealed enhanced spreading and protrusion of Dia1KD-Scarlet cells both atop and below neighbors (Fig. 4F). The wedge-shape associated here with Dia1-deficiency mimicked morphology marking the delamination and protrusion of keratinocytes into the suprabasal layer during stratification (Miroshnikova et al., 2017). Together, the results supported a hypothesis that heterotypic interactions between Dia1-positive and negative cells guided the protrusive activity of Dia1-negative cells towards the suprabasal compartment. The resulting arrangement mimicked the spatial relationship observed *in vivo*.

### Differential Dia1 Expression Drives Cell Sorting

The observed changes in cell morphology implicated Dia1 in modifying cell adhesion and/or cortical mechanics. To query this, we used immunofluorescence to visualize the actin and collagen in vertical cross-section of newly forming adhesions between doublets of CTL or Dia1KD cells 4 and 20 hrs post-plating (Fig. 5A). We observed, at 4 hrs post-plating, that nascent cell-cell contact between CTL cells induced distortion of the underlying collagen; this was not observed at 20 hrs (Fig. 5B). This matrix distortion is accompanied by rapid actin assembly at cell-cell adhesions and the apical surface of the cells (Fig. 5C), consistent with a role of formin-dependent actin assembly supporting cortical tension (Acharya et al., 2017; Bovellan et al., 2014; Harris et al., 2014). Indeed, nascent cell-cell contacts formed by Dia1KD cells did not distort the underlying collagen at either 4 or 20 hrs (Fig. 5A-B) and the apical actin was diminished (Fig. 5C).

**Figure 5.**
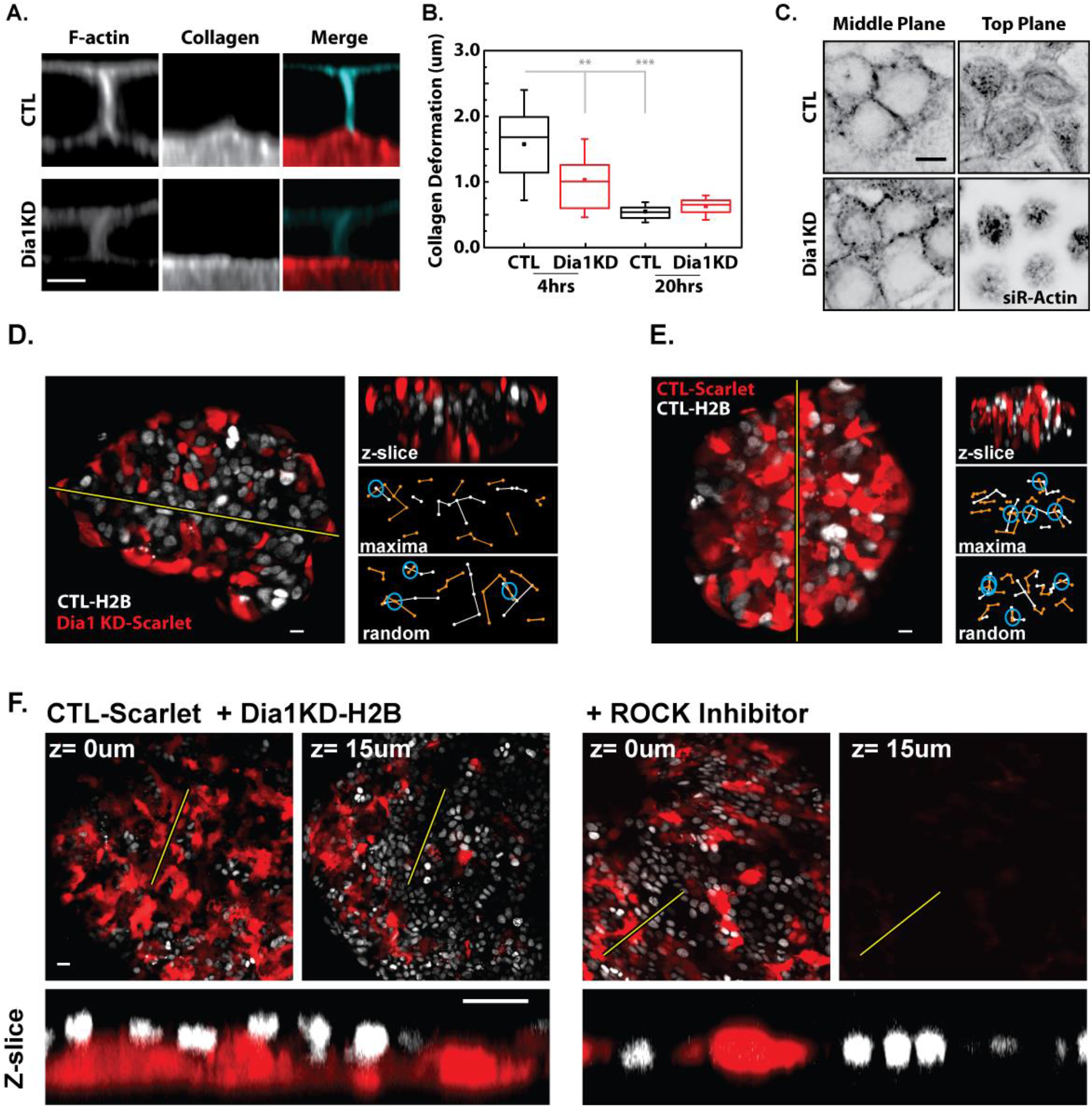
Differential Dia1 Expression Drives ROCK-dependent Separation Into Stratified Layers. A) Orthogonal view of forming junctions, 4hrs post-plating upon labeled collagen. B) Quantification of the vertical collagen deformation observed below forming junctions. Represents 31-36 junctions analyzed at 4hrs, pooled from two experiments. 20hr timepoint represents 13-15 junctions. C) SiR-actin stain of planes taken through the center and top of cells 4hrs post-plating. D) 1000 cell aggregate formed within a hanging drop containing CTL-H2B and Dia1KD-Scarlet cells. Right panels: Z-slice taken along line in left panel and the networks drawn amongst scarlet (orange) and h2b (white) cells connecting local maxima with nearest neighbors. Includes an example of randomized maxima. Blue circles indicate network intersections. E) Control cell aggregate experiment in which two differentially labeled populations of control cells (CTL-Scarlet and CTL-H2B) were mixed and allowed to form aggregates in suspension. Right panel shows cross-section of aggregate taken along yellow line and example intersection analysis as in panel D. F) Larger, 20000 cell aggregates formed with CTL-Scarlet and Dia1KD-H2B (note reversal of labeling compared to panel D), with or without 10μM ROCK inhibitor. Box plots: Boxes= 25-75th percentiles, square= mean, line= median and whiskers= 10-90th percentiles. Scalebars: 5μm in A, 10μm in C, D, and E and 20μm in F. Student’s t-test results are indicated as: p<0.05 (*), p<0.01(**), p<0.0001(***), p>0.05 was considered not significant (ns).

In matrix-free cell sorting assays, differential tension and adhesion can drive the separation of distinct cell populations (Foty and Steinberg, 2005; Maitre et al., 2012). We sought to explore whether sorting of CTL and Dia1KD cells would also occur in a hanging drop assay (Foty and Steinberg, 2005). To form aggregates, CTL cells were mixed 1:1 with Dia1KD cells and suspended from petri dish lids in drops of media. Mixtures totaling 1000, 5000 or 20000 cells were tested. To distinguish populations, cell lines carrying the GFP, nuclear label (CTL-H2B or Dia1KD-H2B) were mixed with those carrying the Scarlet, cytoplasmic labels (CTL-Scarlet or Dia1KD-Scarlet). After 20 hrs, confocal microscopy enabled detection of local maxima in z-slices and networks of nearest neighbors were drawn for each signal. CTL-H2B and Dia1KD-Scarlet cells separated within such aggregates, yielding networks that intersected seldomly compared to those constructed from randomized positions (Fig. 5D). In contrast, there was a high degree of overlap between networks derived from aggregates containing mixtures of CTL-H2B and CTL-Scarlet cells (Fig. 5E). To quantify separation, the number of intersections determined per z-slice from randomized networks was divided by the number of actual, observed intersections. Thus, randomly distributed mixtures would yield a separation index of 1 with higher values representing a greater degree of separation. Analyzing 10 z-slices per aggregate indicated that CTL/Dia1KD mixtures (mean index= 1.40 +/− 0.13 s.e, n=21) displayed significantly (p= 0.04) higher levels of separation than CTL/CTL mixtures (mean index= 1.00+/− 0.05 s.e., n=11).

Though Dia1 coordinates with ROCK-dependent myosin activity to generate tension along intercellular contacts (Acharya et al., 2017), treatment of CTL/Dia1KD mixtures with the Y-27632 ROCK-inhibitor had little impact on the separation index (mean index= 1.41 +/− 0.17 s.e, n=12). However, inspection of three-dimensional structures revealed a key distinction between untreated and inhibitor-treated mixtures. In large aggregates (20000 cells), Dia1-positive and negative cells separated into stratified layers, remarkably, mimicking the arrangement of cells in actual stratified tissue (Fig. 5F). Cells in the presence of ROCK-inhibitor continued to sort but did so horizontally within monolayered aggregates. This could represent local clonal expansion. Alternatively, differential Dia1 expression may drive cell sorting independent of ROCK but rely on ROCK-dependent myosin activity to establish a stratified arrangement. We concluded that when presented with Dia1-negative counterparts, Dia1-positive cells preferentially adhere to each other and that addition of ROCK activity drives this separation into a stratified structure.

### Differential Dia1 Expression Guides Adhesion Rearrangement Following Division

Division in the basal layer figures prominently amongst events which affect cell position in the epidermis (Lechler and Fuchs, 2005; Miroshnikova et al., 2017). In light of the cell sorting data, a reasonable scenario would apply the given role of Dia1 in building tension-bearing actin networks at adherens junctions (Acharya et al., 2017) towards remodeling events associated with division (Higashi et al., 2016). Static analyses also indicate that Dia1 supports lateral adhesion height (Carramusa et al., 2007), suggesting that it may aid in rebuilding lateral adhesions following cytokinesis. To test whether Dia1 expression affected the rate of cell-cell contact expansion, the behavior of nascent cell-cell contacts was observed. CTL or Dia1KD cells, labeled with siR-actin, were imaged immediately after plating (t = 0 m) to capture the formation of cell-cell interfaces along the apical-basal direction. In control cells, the adhesive interface of the lateral membrane domain expanded over the first four hours, increasing to 7 μm in height and increasing the angle formed between cell apices at the top of the contact (Fig. 6A-B, Fig. S4). By contrast, in Dia1KD, the adhesive interface failed to expand over the first 4 hours (Fig. 6A-B). This result represented one of the first attempts to visualize a Dia1-dependent effect on lateral membrane growth kinetics. Reintroducing mDia1 to Dia1KD cells (Dia1-Rescue) also accelerated the accumulation of actin at nascent junctions visualized in the XY plane (Fig. S4). As cells must reallocate adhesive surfaces following division, these data argued in favor of Dia1 expression possibly guiding division-associated rearrangements by favoring an actin-driven expansion of interfaces between Dia1-positive siblings.

**Figure 6.**
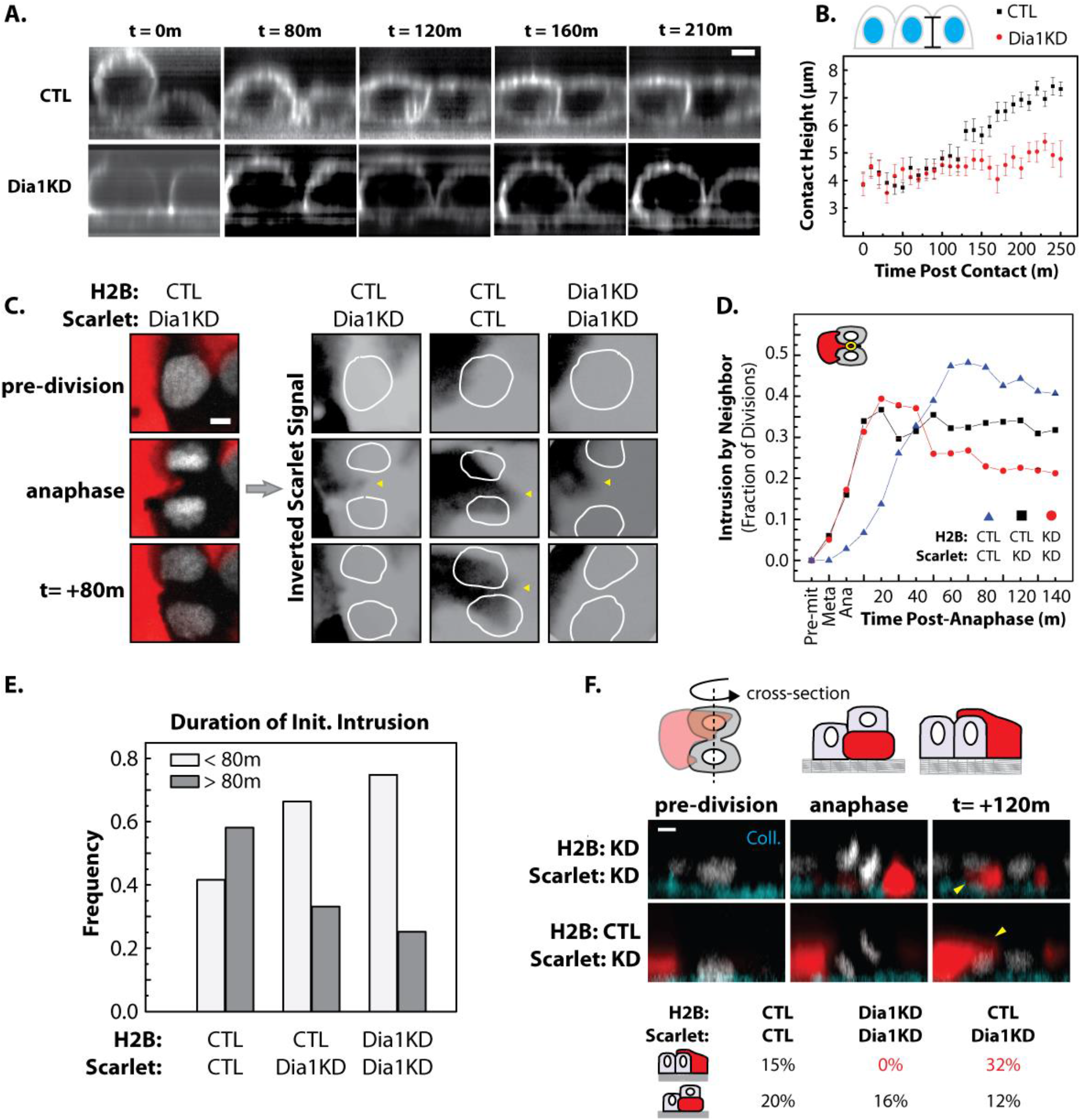
Dia1 Expedites Lateral Adhesion Growth and Favors Reintegration of Dividing, Dia1 Positive Siblings. A) Orthogonal view of cells labeled with siR-actin and tracked from seeding on collagen gels. 5μm scale bar. B) Junction height as a function of time post contact. 20-23 junctions analyzed per condition, +/− s.e. C) Mixed culture demonstrating response of Dia1KD-Scarlet cells to a neighboring CTL-H2B cell division progressing from top to bottom panels. 5μm scale bar. Panels at right demonstrate intrusions (arrows) of Scarlet-labeled cells into dividing, H2B-labeled cells (nuclei/chromosomes outlined in white) for indicated cell mixtures. D) Fraction of divisions in which a neighboring, Scarlet-labeled cell reaches the middle of an H2B-labeled division plane. 20-25 divisions per condition. E) Frequency of short-lived (<80m) and long-lived (>80m) intrusions initiated within 60m of anaphase (analysis ended at 160m). F) Behaviors observed in vertical cross-section, 2hrs post-anaphase. Percent of total divisions in which an H2B-labeled daughter cell packed under or on top of a neighboring Scarlet cell. Scarlet signal was considered basal or suprabasal with respect to H2B labeled cells if it occluded the H2B signal by >50%. 5μm scale bar. A Student’s t-test of data in panel B indicated all differences beyond 120min (except for t=140m) were signficant (p<0.05).

The extent to which adhesion kinetics affected division-associated rearrangement was explored by live imaging of mixed cultures. Juxtaposition of H2B-marked cells with neighbors carrying Scarlet-marked cytoplasm enabled tracking of cell-cell and cell-matrix interfaces during cell division. In all mixtures tested, neighboring cells responded to division by entering the space formerly occupied by the dividing cell, often at the cytokinetic furrow (Fig. 6C). We predicted that, when presented with the option of reintegrating with a Dia1-positive sister or a Dia1-negative neighbor, differential adhesion would favor rapid reintegration of CTL-H2B daughter cells and recession of the Dia1KD-Scarlet neighbor. Conversely, in the absence of differential Dia1 expression, intrusions made by CTL-Scarlet neighbors could become stable, possibly leading to neighbor exchange or stratification. Measuring the frequency with which intruding cells occupied a position between daughter cells as a function of time post-anaphase supported these predictions (Fig. 6D). Measurement of intrusions made within the first hour after anaphase and tracked up to 160m post-anaphase revealed that Dia1 expression favored production of long-lasting intrusions (Fig. 6E). Roughly 60% of intrusions produced by CTL-Scarlet cells lasted >80m. Intrusions produced by Dia1KD-Scarlet cells, in contrast, only exceeded 80m in 25% of attacks made upon Dia1KD-H2B neighbors and ~35% of those made upon CTL-H2B divisions. Thus, juxtaposition of Dia1-positive and negative cells enhanced the probability of Dia1-positive daughter cells reintegrating after division. In an *in vivo* context these behaviors would favor channeling proliferative output into packing the basal layer.

Early in development, when keratinocytes exist as a monolayer, cells often divide in an out-of-plane, oblique manner to begin building stratified structures (Box et al., 2019). In our *in vitro* division analysis, we noted that intrusions aimed initially at the cytokinetic furrow would often expand across one of the daughter nuclei when observed via maximum projections (see Fig. 6C, CTL/CTL panels). This suggested that intrusions were either expanding underneath or spreading atop the daughter cell. To clarify the position of intrusions, orthogonal images were created from confocal z-slices sectioned perpendicular to the division plane (Fig. 6F). At 2hrs post-anaphase, in CTL-H2B/CTL-Scarlet mixtures, 15% of all divisions analyzed resulted in a Scarlet-labeled neighbor expanding over an H2B-marked daughter cell. A roughly equivalent number of divisions (20%) resulted in the Scarlet-labeled cell positioned beneath the daughter, in effect, an oblique division. However, when the neighbor was a Dia1KD-Scarlet cell, 32% of CTL-H2B divisions resulted in one daughter cell packing underneath a Scarlet-marked neighbor. Only 12% assumed the opposite arrangement. In Dia1KD-H2B/Dia1KD-Scarlet mixtures, no divisions resulted in daughters being packed beneath a Scarlet-labeled neighbor, whereas, 16% placed a daughter on top of a neighbor. These behaviors likely account for the static observations made in Figure 4e concerning the position of CTL cells with respect to Dia1KD neighbors. The timing of Dia1 downregulation with respect to stratification is not yet known. However, these data indicate that downregulation of Dia1 by a basal cell would result in delamination and stratification when crowded by dividing, Dia1-positive neighbors.

## Discussion

We show that Dia1 is required to construct a crowded basal layer which, in turn, supports differentiation of stratified epidermal tissue. In its absence, enforced crowding is sufficient for differentiation, indicating a primary role for Dia1 in determining basal layer mechanics. Epidermal tissue relegates YAP activity to undifferentiated basal cells and its inhibition by crowding elicits differentiation (Elbediwy et al., 2016). Accordingly, the present study suggests that Dia1-dependent crowding impinges on this pathway as well as those mediated by Polycomb Repressive Complexes *en route* to supporting differentiation (Fig. 7). We speculate that Dia1 supports crowding by controlling remodeling of cell-cell adhesions required in response to cells added by proliferation. Cells in a monolayer can accommodate an increasingly crowded environment by building taller lateral junctions and occupying a smaller basal footprint or they can avert crowding by delaminating and leaving the monolayer entirely. Our data shows that Dia1 accelerates the growth of lateral cell-cell adhesions, which we surmise favors the option of packing cells into a crowded configuration. Accordingly, Dia1 depletion led to keratinocytes leaving the basal layer before reaching an appropriate density. This ultimately results in stratified tissue marked by aberrant basal layer morphology and a diminished capacity for differentiation.

**Figure 7.**
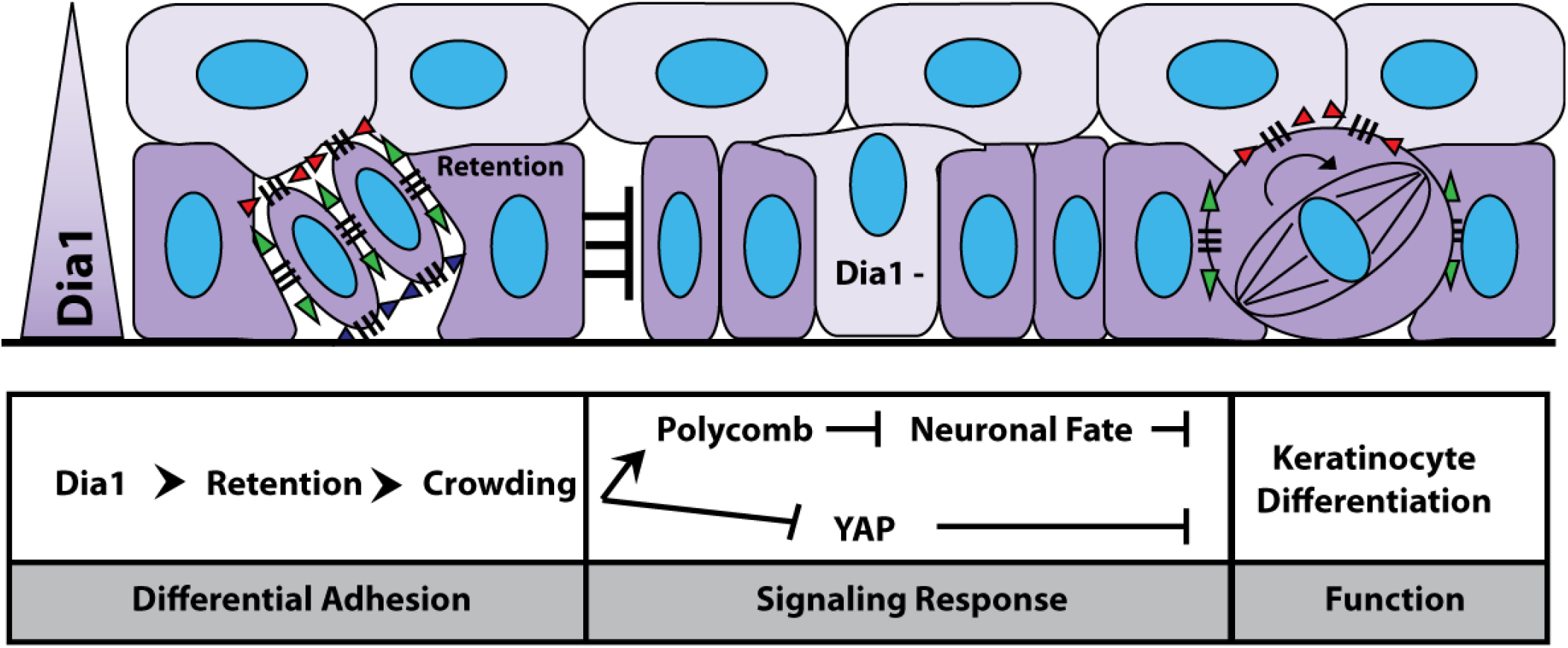
Working Model: The effect of differential Dia1 expression on adhesion supports basal layer packing by favoring reintegration of dividing cells with the basal layer and preventing premature delamination.

This work also has implications for the role of Dia1 suppression in the initiation of tissue stratification. Two dominant theories address the coupling of differentiation to cell position in stratified epithelia. Reoriented division and delamination amongst basal cells both contribute to populating the suprabasal compartment. However, recent intravital imaging reveals that dynamic factors pertaining to adhesion and geometry actively ensure proper execution of these processes (Box et al., 2019; Lough et al., 2019; Mesa et al., 2018; Miroshnikova et al., 2017). Though the present study did not aim to compare delamination with asymmetric division, we posit that preferential adhesions established by Dia1 expression gradients would act in both cases to channel proliferative output into basal layer packing (Fig. 7). Our data show that Dia1-negative cells interact transiently with dividing Dia1-positive cells but these interactions fail to stabilize. *In vivo*, this instability would favor reintegration of dividing, Dia1-expressing cells with the basal layer while driving the Dia1-suppressed cells to delaminate and form a suprabasal layer. A more resolved temporal analysis is required to determine at what timepoint inhibition of Dia1 expression, presumably by miR-198, occurs. However, this study suggests that loss of Dia1 expression amongst basal cells and subsequent delamination, coupled with oriented cell division, could represent a step in the stratification process. More generally, differential formin expression as a mechanism to drive differential adhesion and sorting may widely impact any epithelia in which proliferation must be shunted into populating specific compartments.

Since suprabasal cells lack Dia1 but robustly resist disintegration by exogenous force, we cannot claim that Dia1 is formally required for strong intercellular adhesion in all cases. In the present work, this pointed to adhesion kinetics and cortical mechanics as the behaviors capable of driving cell segregation. Specifically, Dia1 expression accelerates expansion of a lateral contact under tension, capable of distorting the underlying collagen. We postulate that this function favors creation of a columnar shape by expanding the lateral “wall” between two Dia1+ cells at the expense of area dedicated to the apical “roof” or basal “floor”. Thus, Dia1 may specifically populate basal cells because it uniquely provides the polymerization activity necessary to expand an adhesive surface, the lateral membrane, whilst under tension. Previous work has implicated cortical tension in restricting basal layer exit (Miroshnikova et al., 2017). Dia1 is a force-activated nucleator (Zimmermann and Kovar, 2019) and is implicated in cortical mechanics (Bovellan et al., 2014); these suggest particular advantages in controlling cell shape and junction remodeling during division. In the context of rearrangements elicited by basal layer divisions, expansion of lateral Dia1+/+ junctions would outpace expansion of the apical Dia1+/− junction, favoring retention of cells in the basal layer. An effect on cell shape occurs both at the monolayer stage when cells lack suprabasal neighbors as well as in the presence of Dia1-suprabasal cells. Together, one interpretation of these results would suggest that the Dia1 expression gradient functions to cloak suprabasal cells in terms of guiding basal cell shape whilst maintaining mechanically adequate adhesion between the two compartments.

In summary, we have demonstrated that differential formin expression aids in defining the morphology and packing of basal layer epidermal keratinocytes with downstream impacts on differentiation. The study broaches new ground in offering evidence that specific formin expression profiles can guide cell positioning through modulating cell-cell adhesion kinetics. In addition to the implications for other formins and tissues which must channel stem cell proliferation into defined architectures, the present study offers a unique perspective regarding the basic process of cell sorting. One implication of our work is that the adhesion kinetics associated with a formin become particularly meaningful in tissues populated by cells with heterogenous formin expression profiles. Viewing formin expression gradients as determinants of tissue domain architecture represents a novel perspective. Extending this possibility to the entire formin family opens a potentially fruitful avenue for exploration of morphology maintenance in all regenerative epithelia. Towards expanding upon the present work, future studies will begin tackling the potential contribution of Dia1-deficiency to *in vivo* wound healing assays, potentially in the context of skin lesions associated with myelodysplastic diseases (Keerthivasan et al., 2014; Lepelletier et al., 2019).

## Materials and Methods

### Cell Culture and Reagents

The HaCaT line of immortalized keratinocytes was provided by Yu-Ying He (University of Chicago), and was originally developed by the laboratory of Norbert Fusenig (Schoop et al., 1999). HaCaTs were maintained in low calcium DMEM prepared from calcium-free DMEM powder (US Biological,#09800), spiked with 40 μM CaCl_2_, penicillin-streptomycin, L-glutamine and 10% fetal bovine serum depleted of calcium by Chelex-100 (Sigma) treatment. For calcium-induced differentiation experiments, calcium concentration was raised to 2.8 mM for the indicated time span. ROCK-inhibitor (Y-27632, Cayman Chemical) was used at 10 μM.

### Antibodies and Subcloning

The following antibodies were used in this study. Anti-Keratin-10 (ThermoFisher,MS-611-P0) was used at 1:100 for immunohistochemistry. Anti-PCNA (Abcam, AB29) was used at 1:100 for immunohistochemistry. Anti-Dia1 (ProteinTech,20624-1-AP) was used at 1:1000 for western blotting and 1:100 for immunohistochemistry. Anti-GAPDH (Sigma, G9545) was used at 1:2000 for western blotting.

To generate tagged mDia1 constructs, SnapGene software (from GSL Biotech; available at snapgene.com) was used to develop InFusion cloning (Takara) reactions for disabling the miRNA cassette and replacing the GFP in pGIPZ with mScarlet. The mScarlet insert was purchased from IDT as a gene block. Full-length mDia1 (NM_007858), was cloned into pGIPZ-scarlet via PCR amplification and the InFusion cloning system (Takara).

### Experiments Utilizing Coverslip-Supported Collagen Gels and 3D Organotypic Cultures

For experiments conducted on collagen gels, 75 μl of 3.6 mg/ml rat tail collagen I (Corning, #354236) was spread upon activated 25mm diameter coverslips. Activated coverslips were prepared with 3-aminopropyltrimethoxysilane and glutaraldehyde as described in Aratyn-Schaus et al (Aratyn-Schaus et al., 2010). Gel polymerization was induced by exposure to ammonium hydroxide vapor for 4 minutes, followed by neutralization in PBS (pH 7.4). An 8mm (inner diameter) glass cloning ring was placed upon the center of the gel which was, subsequently, incubated overnight at 37°C in culture media prior to receiving cells. Labeling of collagen was accomplished by spiking the mixture,1:50, with AlexaFluor-647-conjugated collagen prepared with a protein labeling kit (ThermoFisher, A20186). Unless otherwise noted, 1.75 × 10^5^ cells were plated per cloning ring.

Production of three dimensional organotypic cultures was based on protocols described by Simpson et al (Simpson et al., 2009) with the following modifications. Collagen I solutions (Enzo, ALX-522-435-0100) of 1.5ml (final concentration 4mg/ml) containing J2 fibroblasts at 4.0 × 10^5^ cells/ml were cast in 3 μm pore filter inserts (Greiner,657630) set in deep 6-well plates (BD, #355467). Fibroblasts were allowed to condition the collagen plug overnight prior to 1.0 × 10^6^ keratinocytes being added per collagen plug. After 2 days of submerged culture, keratinocytes were brought into contact with air and allowed to stratify for the indicated duration.

Both organotypic cultures and keratinocyte cultures grown on coverslip-supported collagen gels were carried out in E-media, described by Simspon et al (Simpson et al., 2009), containing 5% fetal bovine serum.

### Lentiviral Transduction and CRISPR Editing

A pGIPZ construct carrying shRNA targeting nucleotides 1528-1546 of the human DIAPH1 mRNA (NM_005219.4) was purchased from Dharmacon (Cat# V2LHS_43611). The Dia1 shRNA had been previously described by Bovellan et al (Bovellan et al., 2014). pGIPZ constructs were co-transfected with pHR1-8.2-deltaR and a VSV-G pseudotyping plasmid (gifts from M. Rosner) into 293T cells for lentiviral particle production via a second generation system described by Zufferey et al (Zufferey et al., 1997). After concentrating virus using Amicon Ultra-15 30kD centrifugal filters, HaCaTs were incubated 6hrs with concentrated virus in the presence of 4ug/ml polybrene. shRNA expressing cells were then selected for by incubation with puromycin. pGIPZ lentiviral transduction was also utilized for introducing Scarlet-tagged protein. GFP-H2B lentivirus were produced, similarly, but with a pLKO.1-based vector (gift from Elaine Fuchs, Addgene plasmid # 25999; http://n2t.net/addgene:25999; RRID:Addgene_25999). GFP-H2B transduced cells were sorted by the University of Chicago Cytometry and Antibody Technology Core.

Prior to conducting CRISPR editing, parental HaCaT cells were subcloned and tested to ensure that the chosen clone produced differentiated, three dimensional cultures. Alt-R CRISPR-Cas9 crRNA targeting DIAPH1 exon1 (5’-UCU UCU UGU CCC GGG UCC CGG UUU UAG AGC UAU GCU-3’) and Alt-R CRISPR-Cas9 tracrRNA ATTO-550 acquired from IDT were complexed with Cas9 enzyme (GeneArt) and transfected into a suitable HaCaT clone using Lipofectamine CRISPRMAX (ThermoFisher) as per IDT instructions. Via flow cytometry, successfully transfected cells were identified and deposited as single cells into separate wells and allowed to expand. Successful disruption of the DIAPH1 locus in individual clones was determined by western blot and editing at the genomic level was assessed by the CRISPR amplicon sequencing service provided by the Massachusetts General Hospital CCIB DNA Core.

### Western Blotting and qPCR

Cell lysates were collected in Laemmli buffer and sheared with 22 and 25 gauge needles before carrying out amido black protein assays. Western blots were conducted by running equivalent amounts of protein on 4-20% Tris-Glycine SDS/polyacrylamide gradient gels (Lonza) after reduction with 2-mercaptoethanol and heating to 95°C for 5m. Protein was transferred to nitrocellulose, overnight at 4°C in standard Tris-Glycine buffer containing 20% methanol without SDS at 30V. Blots were blocked for 1hr in a 5% milk/ 0.05% Tween-20/ TBS (pH 7.8) solution. Primary antibody incubation occurred overnight at 4°C in blocking buffer followed by washing with TBS/Tween and a 1hr incubation of HRP-conjugated secondary antibody at room temperature. After a final series of TBS/Tween washes, blots were developed using chemiluminescence reagent (Pierce) and film (ThermoScientific, CL-Xposure) development. Blots for assessing shRNA-based knockdowns were done similarly, however, a Licor Odyssey system was used to image fluorescent secondary antibodies and LiCor proprietary blocking buffers and washing instructions were used.

For qPCR, total RNA was collected using NucleoSpin kits (Macherey-Nagel, #740955). First-strand synthesis was carried out with the SuperScript III system (Invitrogen, 18080-051) using an oligo dT primer and 200 ng of total RNA as input. First -strand reactions were diluted 3-fold and 2 μl was used as template in 20 μl reactions containing PrimeTime master mix (IDT,1055772) and PrimeTime pre-designed qPCR primer/probe mixtures from IDT (IVL: Hs.PT.58.39460547; KRT1: Hs.PT.58.24741966; GAPDH: Hs.PT.39a.22214836). qPCR reactions were run on a StepOnePlus system (Applied Biosystems). Relative mRNA levels were calculated using the 2^−ΔΔCt^ method with GAPDH serving as a reference gene.

### Immunohistochemistry,Phalloidin and Nuclei Staining

Three dimensional organotypic cultures were fixed in 10% formaldehyde for 3 days then transferred to 70% ethanol. The University of Chicago Human Tissue Resource Center paraffin-embedded, sectioned and provided hematoxylin/eosin stained slides of the skin cultures. For immunostaining, 5 μm sections were de-paraffinized by baking at 55°C for 1hr followed by sequential bathing in xylene and ethanol. Antigen-retrieval was conducted by holding the sections in a pH 6.0 citrate solution at 95°C for 10m. Sections were then blocked in a 2% BSA/TBS solution and incubated with primary antibodies overnight followed by washing and incubation with secondary antibodies for 1hr. After a final series of TBS washes, sections were mounted in gelvatol.

For staining cultured cells on coverslips, a PBS-buffered, 4% paraformaldehyde solution was applied for 10m (20m for cells on collagen gels). Blocking, permeablization and quenching of paraformaldehyde was carried out by placing coverslips in a 2% BSA/TBS solution containing 0.5% TritonX-100 for 30m. To stain nuclei and the actin cytoskeleton, coverslips were incubated with 5μg/ml Hoechst 33342 (ThermoFisher, H3570) and 13 nM AlexaFluor-labeled phalloidin (ThermoFisher, A12379, A12380, A22287) in 2%BSA/TBS for 30m at 37°C. After washing in TBS, coverslips were mounted in a Tris-buffered glycerol mountant containing N-propyl gallate.

### Forced Cell Crowding

To conduct crowding experiments, 1mm thick sheets of Sylgard184 (Fisher Scientific) with a 10:1 polymer to crosslinker ratio were cured at 70°C overnight. The molds created 12mm diameter, 0.17mm deep wells on the sheet surface. The sheets, as well as, steel washers with an inner diameter and thickness of 25mm and 3mm, respectively, were subjected to UV-ozone cleaning for 2m. Washers were adhered to PDMS sheets, centered around the wells, using PDMS of a 5:1 polymer to crosslinker ratio and baked at 70°C. The 12mm wells were biaxially stretched by clamping the washer down over a 6mm post to increase the well area by 36% or a 4.5mm post to acquire a 18% increase. Stretched and unstretched wells were subjected to UV-ozone cleaning followed by an overnight incubation at 4°C with 200 μg/ml collagen I in 0.017N acetic acid. In unstretched wells, 2.13 × 10^5^ cells were plated in low calcium media. In wells stretched to increase area by 18 and 36%, 2.50 × 10^5^ or 2.90 x10^5^ cells were plated, respectively, such that all conditions received a starting cell number of ~1900 cells per mm^2^. Non-adherent cells were rinsed off at 1hr and cells were incubated a further 16hrs. At that point, stretched wells were released from stretch. Parallel samples were fixed immediately to check cell densities while samples destined for RNA-sequencing were allowed to incubate a further 5h prior to lysis for total RNA purification.

### RNA Sequencing Analysis

Total RNA samples were produced as described above and submitted to the University of Chicago Genomics Facility. mRNA libraries were prepared and subjected to Illumina Hi-Seq 4000 (SR 50bp) sequencing. MultiQC analysis indicated that samples produced between 70-110 million reads with 84-87% being uniquely mapped for each sample. The Illumina BaseSpace software suite was utilized to convert raw reads into transcripts per million (TPM) values for each gene. For forced crowding experiments, two independent experiments were conducted, each containing uncrowded and crowded conditions for Dia1-deficient and Dia1-rescue samples. After converting to fold-change over Dia1-deficient values, average uncrowded, Dia1-rescue values exceeding 2-fold (and exceeding at least 1.5-fold in each replicate) were fed into the PANTHER ontology program to produce unbiased functional annotation of Dia1-induced genes. Average fold change values of Dia1-rescue samples were also used in assessing the response of published gene sets pertaining to various signaling pathways. The reference sets used were as follows suprabasal/differentiated epidermal genes (Joost et al., 2016), skin cornification diseases (Lemke et al., 2014), genes supported by the expression or activity of p63 (Truong et al., 2006), Wnt/β-catenin (Watanabe et al., 2014), Gli/Hh (Lewandowski et al., 2015), Notch (Wang et al., 2014), SRF (Olson and Nordheim, 2010) and YAP/TAZ (Wang et al., 2018). Regarding Gli/Hh, only genes reported by Lewandowski et al (Lewandowski et al., 2015) as reduced 2-fold by SHH knockdown with FDR<0.05 were queried. Since a different level of crowding was reached in each experiment, the crowded/uncrowded ratios are reported separately for each experiment rather than as averages. The enrichment of neuronal and polycomb targets amongst genes both reduced greater than 2-fold by re-expressing Dia1 on average and 1.25-fold upon 36% crowding was identified using GSEA software (Broad Institute).

These experiments aimed to identify early indications of differentiation and, in turn, low abundance genes were not filtered out of datasets. As such, trends within gene sets (e.g. differentiation genes, Dia1-responsive genes, etc) were deemed more meaningful than changes to individual genes. Specific genes listed in Figure 3 are provided as examples but require more rigorous testing to achieve statistical significance.

### Hanging Drop Aggregation Assay

Cells marked with GFP-H2B and cytoplasmic mScarlet were mixed 1:1 following trypsinization, then, suspended in 20 μl droplets of high calcium (2.8mM) media on petri dish lids overhanging a pool of PBS. The aggregates, containing between 1.0 to 20 × 10^3^ cells were cultured overnight at 37°C in a cell culture incubator. Aggregates were then fixed with paraformaldehyde and transferred to a coverslip with PDMS applied to edges as a spacer. After rinsing with PBS, gelvatol was used to mount aggregates prior to imaging aggregates in a series of 0.25 μm Z-sections.

### Microscopy and Live Cell Imaging

Confocal microscopy of fixed and live samples was conducted on an inverted microscope (Ti-E; Nikon) with a confocal scan head (CSUX; Yokogawa Electric Corporation) and laser-merge module housing 491-, 561-, and 642-nm laser lines (Spectral Applied Research), a stage controller (Prior), and a cooled CCD (HQ2-Roper Scientific) or CMOS (Zyla-Andor) camera. Images were acquired using 20× 0.75 NA Plan Fluor multi-immersion, 40× 1.15 NA Plan Apochromat water immersion extra-long working distance and 60x 1.2 NA Plan Apochromat water immersion objectives (all from Nikon).Live-cell imaging utilized a stage incubator for temperature, humidity, and CO_2_ control (Chamlide TC and FC-5N; Quorum Technologies). The stage adapter, stage cover, and objective were maintained at 37°C, whereas humidified 5% CO_2_ air was maintained at 50°C at its source to prevent condensation within its tubing. Unless otherwise noted, serial z-sections were done at 0.25 μm steps for fixed samples and 0.5 μm steps for live samples. For live, lateral membrane growth studies, cells were pre-incubated with 50nM siR-actin (Cytoskeleton) for 1hr prior to trypsinization and immediate transfer to an imaging chamber. A 50nM siR-actin concentration was maintained in the imaging media.

### Image Analysis and Processing

ImageJ and MatLab were used for image analysis. To make basal nuclei more visible in images of living H2B-labeled 3D cultures the following filtering technique was applied to average projections of the planes spanning 10 μm up from the lowest in focus nuclear signal. The original image was modified by applying a sigma= 40 pixel median filter. The original image was then divided by the transformed image to remove uneven background signal. For studies concerning positional relationships, H2B-labeled cells were considered to have a suprabasal neighbor if Scarlet signal from a neighbor occluded the nuclear signal by >50%. For studies utilizing only H2B-labeled cells, cells were considered suprabasal if their H2B signal occluded >50% of an underlying H2B signal or could be seen to overlie 2 or more H2B-labeled nuclei. Collagen deformation below forming junctions was taken to be the distance between the line connecting the lowest observed collagen positions between two adjacent cells and the highest position marked at forming junctions.

Measurement of cell sorting was conducted by taking serial z-slices through cell aggregates, then, reconstructing 10 orthogonal, radially configured slices per aggregate using ImageJ. Positions of local maxima for Scarlet (noise=300 px) and GFP- H2B (noise=60 px) signals were ascertained. MatLab was utilized to determine and connect via a line each maximum to the nearest neighbor bearing the same signal. For each cell aggregate the number of intersections and maxima detected amongst the 10 slices was summed and used to calculate the average number of intersections per maxima. Matlab was used to rearrange actual maxima positions to randomized positions within the spatial constraints corresponding to each slice. Dividing the intersections per maxima value for randomized maxima by that acquired from actual observations produced, for each aggregate, a cell separation index score. Higher scores indicated less interaction between networks and a higher degree of cell sorting.

To quantify the extent to which neighboring Scarlet labeled cells invaded the space of dividing, H2B-labeled cells, the Scarlet signal was turned into a binary mask. At each timepoint after anaphase, the presence of this mask within a 5 μm circle centered between the chromosomes of what would become two daughter cells was counted as an intrusion. The circular region of interest was centered upon pre-mitotic and metaphase nuclei to establish a baseline level of intrusion.

### Statistics

Where indicated, independent two-sample Student’s t tests were utilized to calculate statistical significance with p-values < 0.05 indicative of a significant difference being detected between means.

## Supplemental Material

Additional figures pertaining to culture histology and immunostaining, genomic and transcript sequencing, and proliferation are contained in the supplemental material.

## Acknowledgements

The above work was supported by the following funding entities: M.L.G. acknowledges funding from NIH RO1 GM104032. RMH acknowledges funding via an NIH National Research Service Award (NIGMS:1F32GM117928-01). The authors declare no competing financial interests.

## Author Contributions

Robert Harmon conceptualized and wrote the manuscript, in addition, to devising, performing and analyzing experiments. John Devany aided in quantitative analysis, concept development and manuscript preparation. Margaret Gardel aided in manuscript preparation, conceptualization, experiment planning and data interpretation.

