## Supplemental Figures for "Dia1 Coordinates Differentiation and Cell Sorting in a Stratified Epithelium"

**Supplemental Figure Legends:**

**Supplemental Figure 1. Disruption of DIAPH1 gene expression and example of ingressions found in DIAPH1-deficient cultures.** A) Western blot analysis of HaCaT cells transduced with non-targeting (NT) and DIAPH1-directed (Dia1) shRNA via lentivirus, cultured in low or high calcium media for 20hrs. Relative abundance indicated with respect to control samples. B) Western blot analysis of control cells (CTL) and those carrying a disrupted DIAPH1 locus (CRISPR) at two different passages. Relative abundance to controls is indicated. C) NGS analysis of genomic DNA flanking CRISPR-editing site and variant frequency. D) Example of DIAPH1-deficient tissue growth into the underlying collagen substrate. Hematoxylin and eosin stain of cross-sectioned, paraffin-embedded tissue fixed 10 days after raising to an air-liquid interface. Scale bar= 20 microns. Yellow arrow indicates basal keratinocyte layer. Red arrows indicate distinct ingressions.

**Supplemental Figure 2. Organotypic cultures treated with shRNA, ectopic mDia1 constructs and ancillary RNA-seq data.** A) Hematoxylin and eosin (H&E) staining and keratin-10 (KRT10) staining of organotypic cultures transduced with control or Dia1-targeted shRNA. Counterstained with Hoechst for DNA. Dotted line indicates keratinocyte/collagen boundary. Scale bars = 20µm. B) Map of mDia1 construct introduced to CRISPR-edited Dia1 knockdown (Dia1KD) cells via lentivirus. Numerals are referenced in subsequent panels. FL= full length. C and D) Western blot analysis of ectopic constructs blotted with anti-DIAPH1 (C) or anti-Cherry (D). Residual Scarlet signal indicated by arrowhead. Irrelevant lanes of western blot lanes were dimmed for clarity. E) RNA-seq analysis of DIAPH1 transcripts in CRISPR-edited Dia1KD cells. All detected transcripts contained deletions in the first exon. F) RNA-seq derived quantities of other mammalian formin genes. Values in parentheses represent average transcripts per million (tpm) value in Dia1KD cells from two experiments. Levels in Dia1-Rescue cells (ii) were normalized to this value and are indicated in the heatmap. DAAM2 was not detected in Dia1KD cells, thus, the average tpm for rescue cells is given.

**Supplemental Figure 3. Response of known gene sets to re-expression of Dia1.** RNA sequencing results tracking the expression of transcripts known to be upregulated by the indicated signaling pathways, comparing Dia1-Rescue cells relative to Dia1KD-Scarlet counterparts . Studies identifying the plotted gene sets are listed in the Methods section. SPRR (small proline rich repeat proteins) are known markers of keratinocyte differentiation. All SPRR genes listed in the NCBI Gene database with non-zero expression values in both the Dia1-deficient and Dia1-rescue lines are plotted. Data represent the average fold change value of individual genes derived from two independent experiments, corresponding to the uncrowded conditions described in Figure 3. Average values were converted to Log2 form for presentation.

**Supplemental Figure 4. Effects of Dia1 expression on proliferation, intercellular contact angles and junctional actin.** A) Hoechst stained nuclei 24hrs post plating on collagen gels. Arrowheads mark mitotic cells. Cell densities indicated. Percent of nuclei with mitotic morpohology was determined from 1131 and 1136 CTL and Dia1KD cells, respectively. s.d.= standard deviation. field = 27500 μm2 .  B) Angle formed between cell apices and the top of developing intercellular junctions labeled via siR-actin and imaged live after contact initiation. 20-22 junctions analyzed per condition. C) Phalloidin staining of F-actin in fixed Dia1KD and Dia1-Rescue cells 1hr after plating on collagen gels, imaged 2.5μm above the substrate.  Scalebars = 20μm for A, 10μm for C. Box plots= 25-75th percentile, line= median, square= mean, whiskers = 5-95th percentile and circles mark measurements for individual cells.

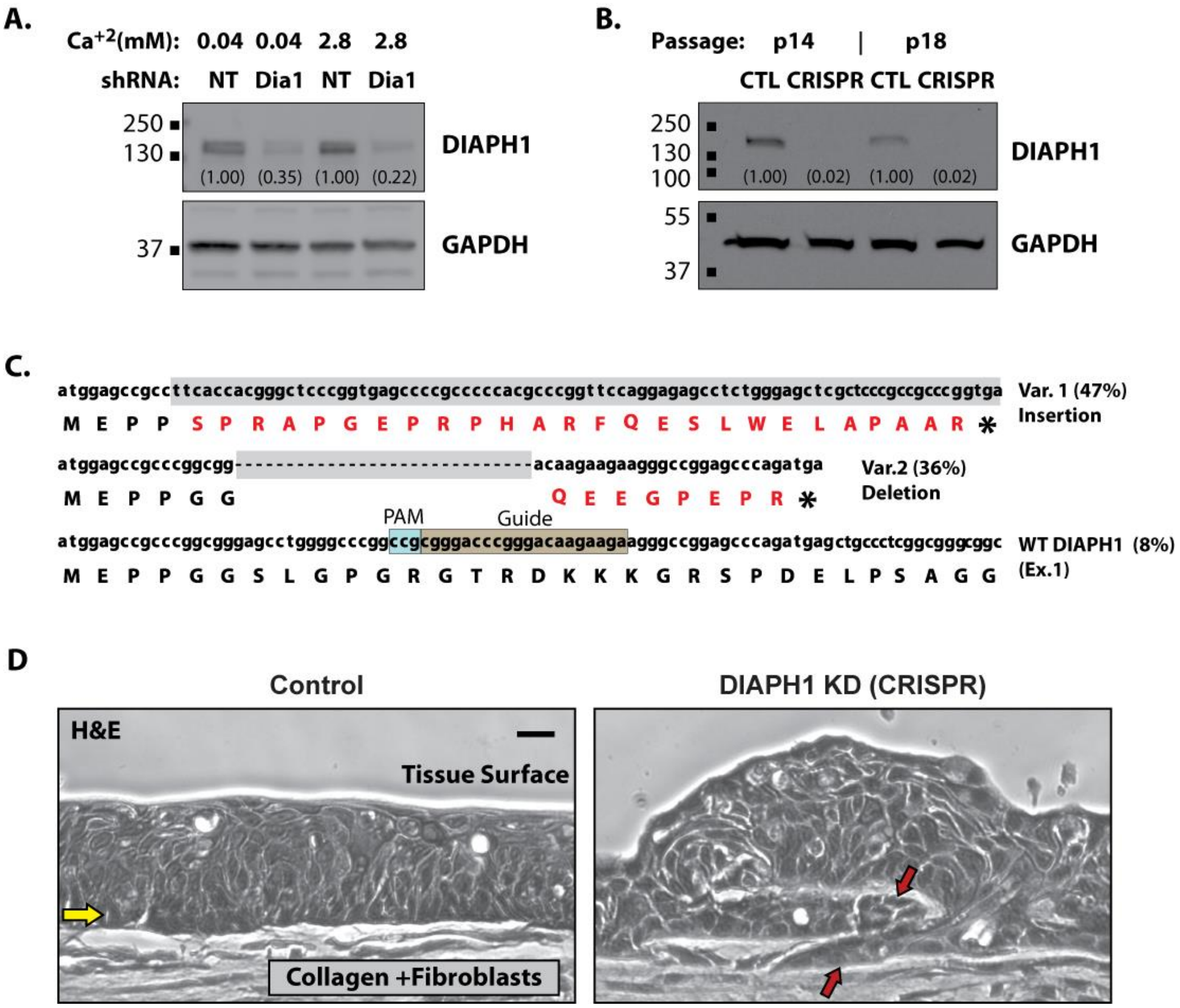


Supplemental Figure 1

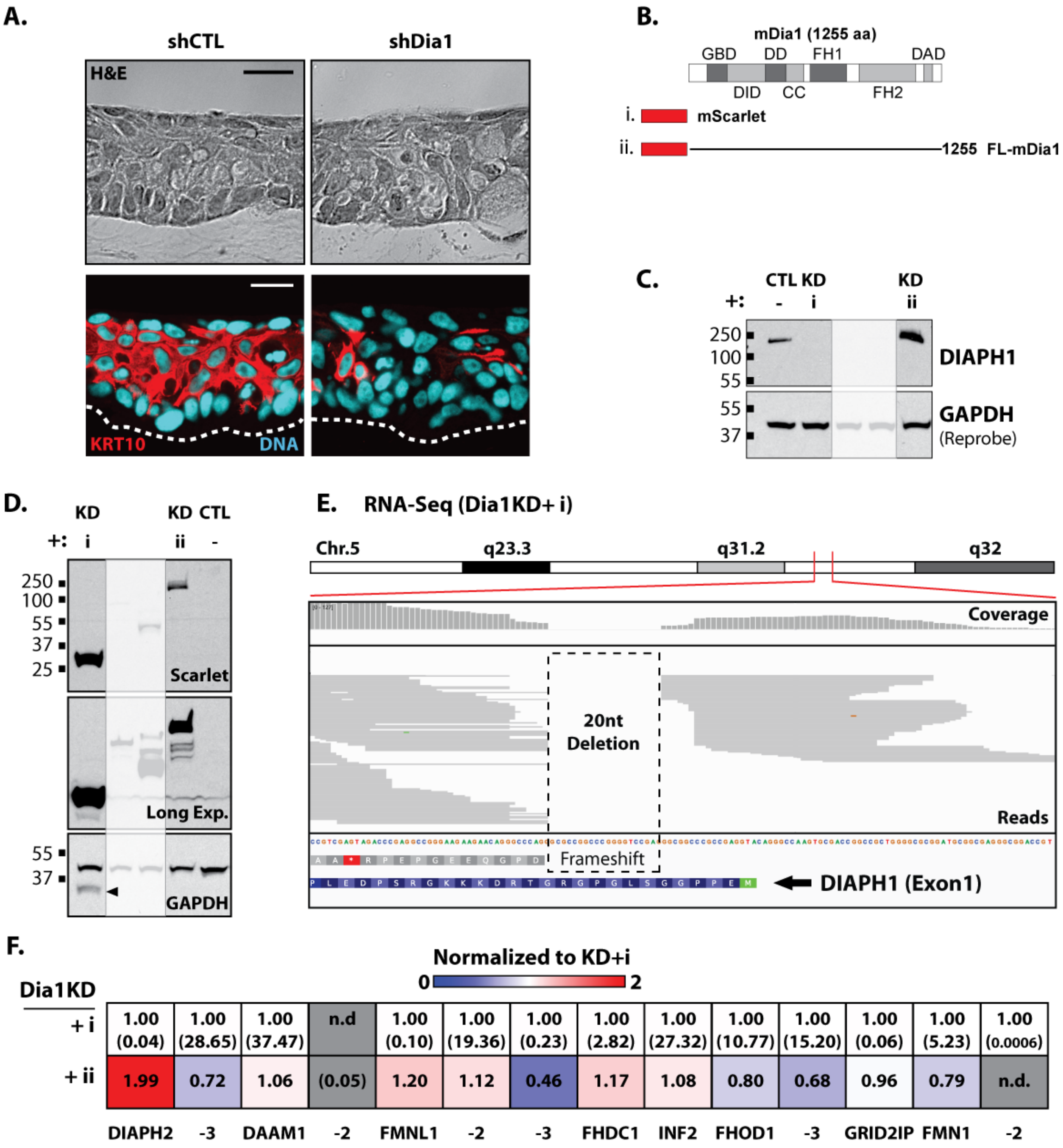


Supplemental Figure 2

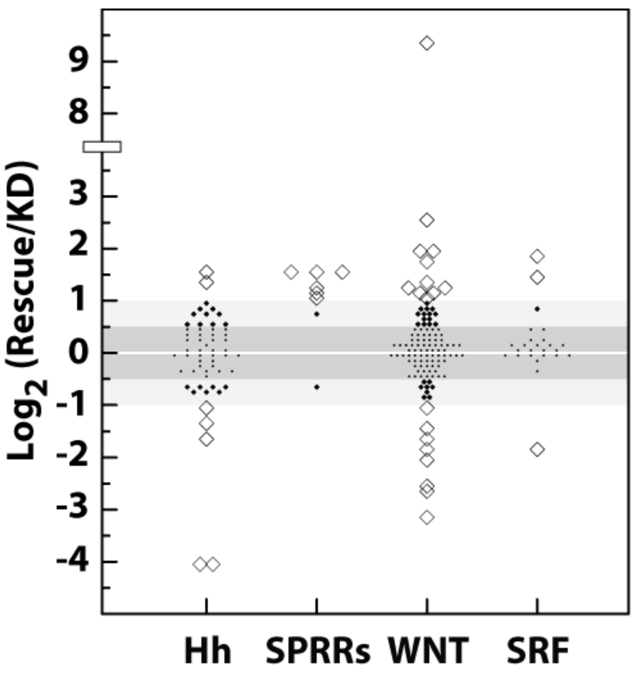


Supplemental Figure 3

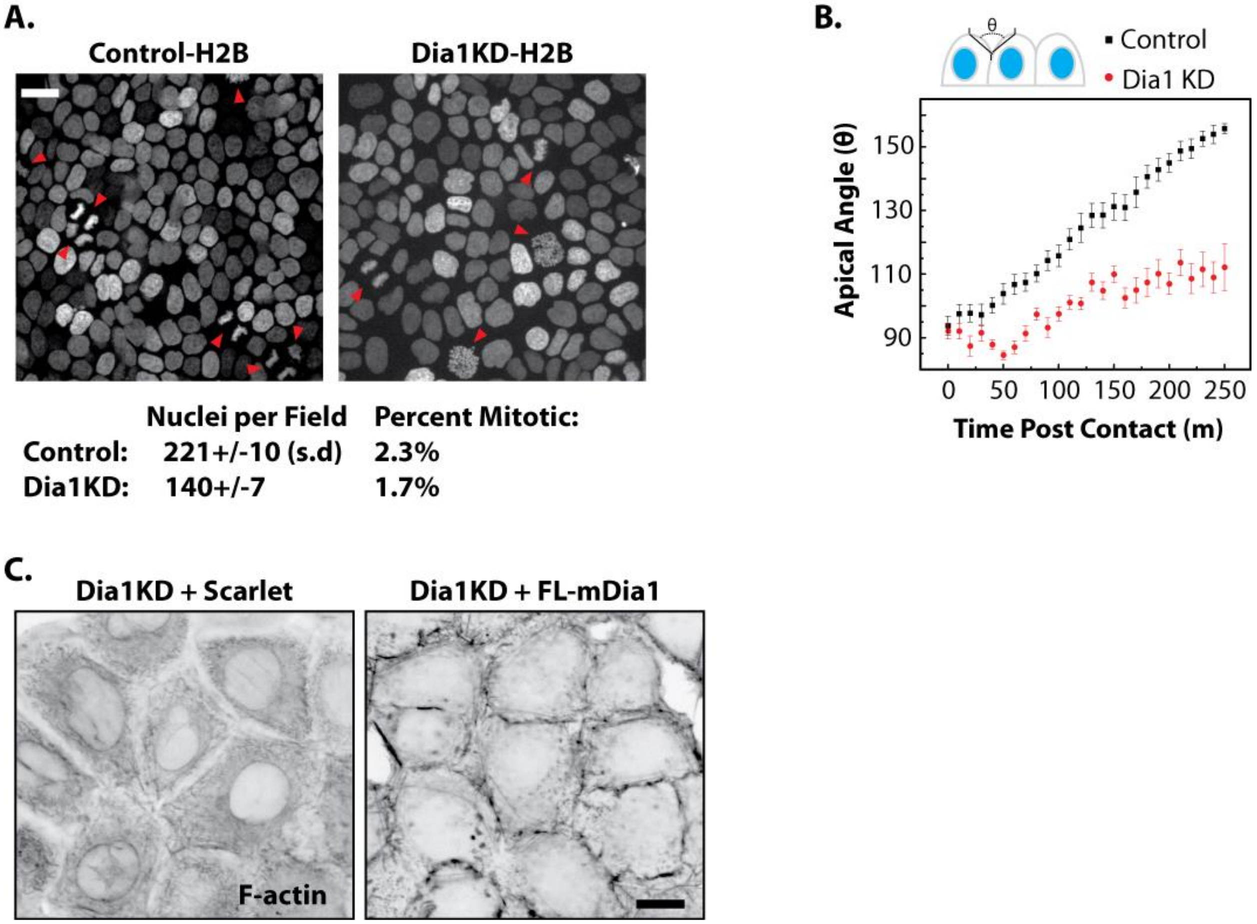


Supplemental Figure 4
